# Impact of water models on structure and dynamics of enzyme tunnels

**DOI:** 10.1101/2023.04.19.537534

**Authors:** Aaftaab Sethi, Nikhil Agrawal, Jan Brezovsky

## Abstract

Protein hydration plays a vital role in many biological functions and molecular simulations are frequently used to study the effect of protein hydration at the atomic level. However, the accuracy of these simulations has often been highly sensitive to the water model used, a phenomenon best known in the case of intrinsically disordered proteins. In the present study, we investigated the extent to which the choice of water model alters the behavior of complex networks of transport tunnels. Tunnels are essential because they allow substrates and products to access and exit the active sites of enzymes that are otherwise deeply embedded within the protein structure. The ability of these tunnels to regulate access directly affects enzyme efficiency and selectivity, making their study crucial for understanding enzyme function and inhibition at a mechanistic level. By performing all-atom molecular dynamics simulations of the wild-type haloalkane dehalogenase LinBWT and its two variants, LinB32 and LinB86, with synthetically engineered tunnel networks in TIP3P and OPC water models, we investigated the effects of these models on the overall tunnel topology. We also analyzed the properties of the main tunnels, such as their conformation, bottleneck dimensions, sampling efficiency, and duration of the tunnel opening. Our data demonstrate that all three proteins exhibited similar conformational behavior in both water models and differed in the geometrical characteristics of their auxiliary tunnels, in line with experimental observations. Interestingly, the results indicate that the stability of the open tunnels is sensitive to the water model and the system under question. Our findings suggest that the 3-point TIP3P model can provide comparable inference on the overall topology of the networks of primary tunnels and their geometry, and thus may be a desirable choice when computational resources are limited or when compatibility issues impede usage of OPC with certain protein force fields. However, when a more thorough investigation is performed, such as the calculation of ligand unbinding rates via such tunnel networks, where precision and intricate details are paramount, the more costly 4-point OPC model would be more suited.

**Graphical Abstract:** 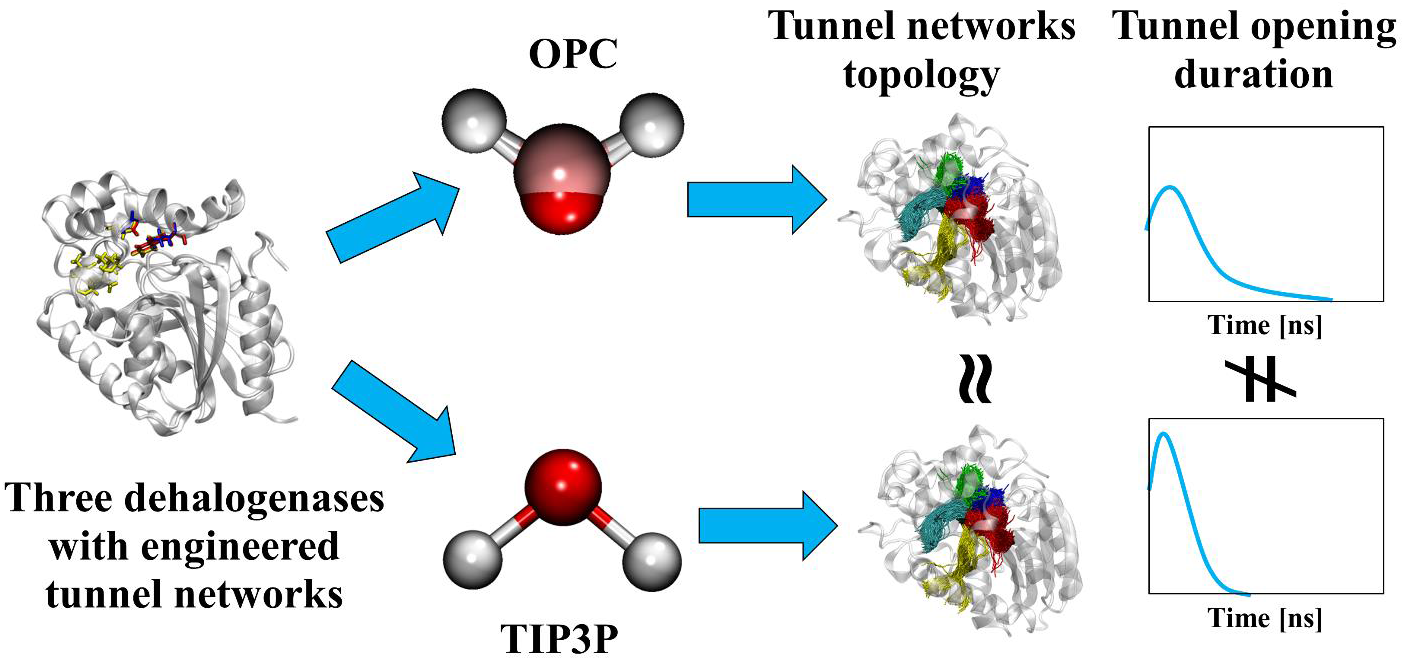

## 1. INTRODUCTION

Water is indispensable in determining the structure, stability, dynamics, and function of proteins.^1^ It plays a critical role in various biological processes such as protein folding and enzymatic catalysis.^2,3^ Interfacial/bound and internal/buried water molecules can affect protein-protein association, protein-ligand interaction,^4,5^ protein thermal stability, and peptide/ligand binding affinity.^6,7^ Furthermore, proteins also perturb the dynamics of water molecules. The surface of proteins is highly heterogeneous due to geometrical disorder and the different energetics of the local H-bonds between water and protein sites.^8,9^ This can lead to increased residence times of some water molecules in the vicinity of protein surfaces. It has been estimated that the residence time of such dampened water molecules near the protein surface could be in the range of nanoseconds to microseconds.^10,11^ However, the residence time of water molecules buried inside the protein structure could reach up to milliseconds.^1,12^ In addition, the proximity to proteins can also affect the re-orientational and translational behavior of water molecules, with NMR studies observing that water reorientation and translation near globular proteins is 3−5 times slower than in bulk.^9,13,14^

All-atom molecular dynamics (MD) simulations have been widely employed to investigate protein-water interactions and have provided insight into water properties and their effects on biomolecules.^15,16^ The accuracy of simulations of protein immersed in explicit water molecules depends on the choice of the protein force field and water model used.^17–19^ While there is a large number of water models, the most commonly applied are Single Point Charge (SPC),^20^ Transferable Intermolecular Potential (TIP),^21^ and Optimal Point Charge (OPC).^22^ The SPC family includes three-point charges placed on the nuclei and a single 12-6 LJ term centered on the oxygen or hydrogen atoms.^20^ The TIP family consists of 3-, 4-, and 5-point water models named as TIP3P, TIP4P, and TIP5P, respectively.^23^ Finally, the OPC family includes 3- and 4-point water models OPC3 and OPC.^24^ These water models, while imperfect, provide a good compromise between accuracy and efficient calculations. For example, the kinetic properties of TIP3P, such as self-diffusivity and viscosity, do not agree well with the experimental observations, but the thermodynamic properties are considered to be a reasonable approximation.^1,25^ Historically, most of these water models have been developed in conjunction with a particular protein force field, such as SPC with GROMOS,^26^ TIP3P with AMBER and CHARMM,^27,28^ and TIP4P with OPLS force fields.^29^ Due to its less tight coupling to any particular water models, AMBER force fields have been shown to work with TIP4P, TIP5P, and OPC water models.^30,31^

To date, many studies have investigated discrepancies that arise from using different explicit water models. For example, Anandakrishnan *et al*. used TIP3P and TIP4P/EW models to investigate the protein folding landscape of the mini-protein CLN025 and found almost order-of-magnitude differences in the models’ ability to predict the fraction of unfolded states. The origin of this discrepancy was traced to the water-water electrostatic interactions.^19^ Gupta *et al*. studied temperature-dependent glass transition of Trp-cage mini-protein using mTIP3P, TIP4P, and TIP4PEw models and found that the TIP3P model resulted in a decreased number of water molecules packed in the first hydration shell of the Trp-cage surface.^32^ Importantly, conformational ensembles of intrinsically disordered proteins simulated with different water models were shown to differ, often being too compact in comparison to various experimental measurements, which could primarily be ascribed to weaker dispersion interactions.^33–37^ Considering the impact of models on the binding of water molecules to proteins, Fadda *et al*. performed simulations of the Concanavalin A protein in free and ligand-bound forms in TIP3P, TIP4P, and TIP5P models. The authors demonstrated that the binding energies of the isolated water molecule are highly sensitive to the model.^38^ Finally, a study on water diffusion through aquaporin AQP1 by Gonzalez *et al*. compared TIP3P, OPC, and TIP4P/2005 models.^39^They showed that OPC and TIP4P/2005 were able to reproduce protein-water interactions in good agreement with experimental data, while the application of TIP3P model resulted in overestimated diffusibility of water molecules. A similar observation was recently made by Thirunavukarasu *et al*. also for water transport via tunnels of three distinct globular enzymes.^40^ Overall, these studies suggest that the accuracy of a simulation is rather sensitive to the water models used, which were initially developed with relatively stable globular proteins in mind. This is particularly relevant when considering less ordered protein fragments as well as the interactions and behavior of water molecules buried within the protein.

From this perspective, it is pertinent to consider another structural element of proteins, “tunnels”. These empty spaces connect the surrounding solvent to the buried functional site within certain enzymes.^41–43^ Tunnels are often equipped with molecular gates, switching between closed and open states, making them transient.^44^ The dynamics and structure of tunnels play a critical role in the entry of substrates into the active site and the release of products into the bulk solvent.^44,45^ Another important role of tunnels is to act as a selective filter, allowing only the cognate molecules of substrates or solvents to enter the active site.^46–48^ Interestingly, the sensitivity of the tunnels to solvent composition was demonstrated by Stepankova *et al*., they showed significant alterations of tunnel entrances of three different dehalogenases when exposed to three different aqueous solutions containing water-miscible organic co-solvents.^49^ Since the tunnels are critical for enzyme function, can be rationally engineered to provide improved biocatalysts,^41,45^ and represent attractive drug targets,^50^ it is critical to understand the role of water models on the nature of the tunnel networks observed in the simulated conformational ensembles.

In the present study, we investigated the effect of two different water models on the structure and dynamics of complex tunnel networks in enzymes. For this purpose, we chose the TIP3P water model because of its widespread use,^51^ and the OPC model, which is one of the best models for reproducing bulk water properties.^24^ As a model system with well-understood and diverse tunnel networks, we have used members of the haloalkane dehalogenase (HLDs) family, LinBWT and its two engineered variants, a single-point mutant LinB32 and a four-point mutant LinB86 **(Figure 1)**.^45,52,53^ Structurally, these enzymes belong to the α/β-hydrolase superfamily, characterized by their main and cap domains, with the buried active site at their interface.^54^ HLDs have a long-standing history as a model system for the catalytic action of enzymes with buried active sites due to the wealth of data collected on the transport component of their function.^55^ This includes identifying rate-determining steps,^45,52,53^ numerous crystal structures, and in-depth studies of the effect of mutations in the tunnel networks of these enzymes.^41^ The three HLD model systems used here have different networks of primary and auxiliary tunnels. In LinBWT, the main tunnel (P1) is prominently open, but in LinB32 and LinB86 variants, the P1 tunnel is blocked by the L177W mutation, which hinders substrate and product transport.^45,53^ In addition, Trp177 forms a hydrogen bond to Asp147 which sets up the cap domain gate of the P1 tunnel.^52^ In LinB86, three additional mutations (W140A+F143L+I211L) were introduced into the auxiliary access tunnel P3, resulting in the activation of P3 tunnel and also facilitating enhanced flexibility of the cap domain gate of the P1 tunnel.^45,52^ These different tunnel networks allowed us to analyze the extent to which the water model used affects our ability to detect relevant tunnels and the duration for which the tunnel remains open. Additionally, we analyzed how fast the tunnels were detected, the number of tunnels detected and probed their bottleneck radius, examining any significant differences between the two models.

**Figure 1:**
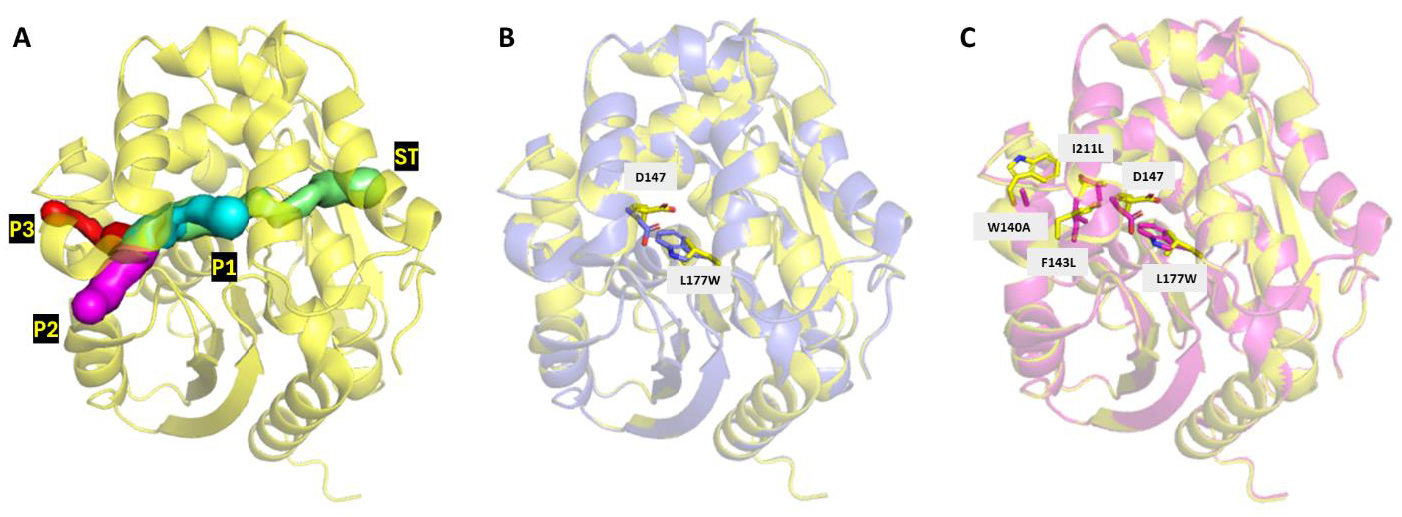
Illustration of the tunnel networks analyzed in the study and residue differences between the three systems. **A**) The P1 (cyan), P2 (magenta), P3 (red), and ST (green) tunnels in the structure of LinBWT (PDB ID: 1MJ5) are depicted in yellow. **B**) The L177W mutation in LinB32 (PDB ID: 4WDQ) in purple aligned onto LinBWT (yellow). This mutation leads to the closure of the P1 tunnel in LinB32 when a hydrogen bond forms between W177 and D147. **C**) The four mutations (L177W, W140A, F143L, and I211L) present in LinB86 (PDB ID: 5LKA) in magenta, aligned onto LinBWT (yellow). The three mutations except L177W are responsible for P3 tunnel activation.

## 2. METHODS

To study the effect of different water models on tunnel dynamics, we have performed simulations for LinBWT, LinB32, and LinB86 enzymes in TIP3P and OPC water models. Initial structures of LinBWT (PDB ID: 1MJ5),^56^ LinB32 (PDB ID: 4WDQ),^45^ and LinB86 (PDB ID: 5LKA)^45^ were prepared for simulations as previously described.^45^ Water molecules were initially placed around the protein structures in three steps. In the first step, 3D-RISM theory was used to add initial water molecules to protein structures according to the Placevent algorithm.^57,58^ In the second step, 3D-RISM predicted water molecules were merged with crystallographic water molecules and in the last step, water molecules that had their oxygen atoms within 2 Å (1 Å = 0.1 nm) of protein atoms were removed. Truncated octahedral water boxes (TIP3P and OPC) were added to the distance of 10 Å from any atom in the respective systems, and ions (Na^+^ and Cl^-^) were added to make a final concentration of 0.1 M using the *tleap* module of AMBER 18.^59^ Hydrogen mass repartitioning (HMR) was performed using the *parmed* module of AMBER 18 to enable a 4 fs timestep.^60^

The system energy was minimized using the *pmemd* module of AMBER 18, by using 100 of the steepest descent steps, followed by 400 conjugate gradient steps within five rounds of decreasing harmonic restrain as follows 500 kcal.mol^−1^.Å^−2^ applied on all heavy atoms of protein, and then 500, 125, 25 and 0 kcal.mol^−1^.Å^−2^ on backbone atoms only. The equilibration of the systems was performed in four stages. In the first round, 20 ps NVT simulations were performed to gradually heat the systems from 0 K to 200 K, followed by 1 ns NVT simulations to reach the target temperature of 310 K. In these two stages, harmonic restraints of 5.0 kcal.mol^−1^.Å^−2^ were applied on the heavy atoms of proteins, and the temperature was controlled using a Langevin thermostat^61^ with a collision frequency of 2.0 ps^-1^. The third stage consisted of 1 ns NPT simulations performed at 310 K using the Langevin thermostat at a constant pressure of 1.0 bar using a Monte Carlo barostat while keeping restraints of 5.0 kcal.mol^−1^.Å^−2^ on backbone atoms only. The last stage employed the same settings as the third but without any positional restraints.

Finally, unrestrained NPT simulations were run for all investigated systems for 500 ns, saving the data to trajectories using 20 ps intervals. All simulations employed the SHAKE algorithm^62^ to fix bonds that contained hydrogens and periodic boundary conditions, with the nonbonded cutoff of 8 Å. The simulations were performed using the *pmemd*.*cuda* module of AMBER 18 using the ff14SB force field.^63^ Three replicated simulations were conducted for LinBWT in TIP3P and OPC water models. In contrast, more simulations were performed to obtain four replicates (two open and two closed forms) of LinB32 and LinB86 in TIP3P and OPC water models **(Table S1)**. The initial 100 ns of simulations were dedicated to water and structural equilibration, leaving 400 ns (20,000 frames) of production simulations for further analyses per replicate.

The dynamic behavior of tunnels was analyzed using the divide and conquer approach,^64^ utilizing CAVER 3.0.^65^ Briefly, 20000 pdbs were generated from each run corresponding to 20,000 frames. These were sliced into 10 batches of 2000 pdbs each. The Caver configuration file was further prepared: the starting point was defined by the center of mass of Trp109, His272, and Asn38. Tunnels were investigated in each trajectory snapshot using a probe radius of 0.7 Å, shell radius of 3 Å, shell depth of 4 Å and a clustering threshold of 4.5 Å. Next, a filtering criteria was applied to the Caver run. Wherein, only those tunnels were retained which were seen in more than 5% of the frames. Afterward, TransportTools was utilized to merge the tunnels.^64,66^ The ward clustering method was applied to cluster the tunnels with a clustering cutoff of 1 Å. The minimum tunnel radius for clustering was set to 0.7 Å and the minimum length tunnel length to 5 Å. Further, the clusters generated by TransportTools were converted to Caver output format using Python script (tt_convert_to_caver.py).^64^ Finally, a comparative analysis was performed using TransportTools to generate individual supercluster statistics. The comparative group definition involved six systems LinBWT OPC & TIP3P, LinB32 OPC & TIP3P, and LinB86 OPC & TIP3P. Here too, the ward clustering method was applied to cluster the tunnels with a clustering cutoff of 1 Å. The minimum tunnel radius for clustering was set to 0.7 Å and the minimum length tunnel length to 5 Å.

Root-mean-square deviation (RMSD), the radius of gyration, was calculated for residue numbered 11-295 backbone atoms (N, CA, C), as the residues 1-10 are loop residues. Residue-wise root mean square fluctuation (RMSF) was calculated using backbone atoms (N, CA, C) of residues 1-295. Radial distribution function (RDF) was calculated for water (O, H1, H2 atoms), around Asp108 (CG atom), with a bin spacing of 0.1 Å, and a maximum bin value of 15 Å. All these analyses were performed using the *cpptraj* module of AMBER 18.^67^

## 3. RESULTS AND DISCUSSION

### 3.1 Comparison of Geometrical Characteristics of Main Tunnels and Their Response to Engineering between OPC and TIP3P

The production phases of all selected simulations exhibited a very stable structure concerning their initial conformations as well as very consistent RMSF profiles between the two water models used **(Figures S1-S3)**. Finally, the relative solvation levels of the transport tunnels showed a high concordance between the simulations in both water models based on their RDFs (**Figure S4**). Overall, these analyses confirmed the absence of major global differences between conformations of analyzed enzymes and their overall solvation when simulated in TIP3P and OPC, allowing us to focus on the tunnel networks. A tunnel works as a dynamic filter that plays an essential role in enzyme activity. Opening the tunnel facilitates the entry and exit of substrates, products, ions and solvent molecules in or out of the enzyme.^50^ To understand the effect of water models on the presence of the key parts of the tunnel network, we calculated how often the tunnels were identified with a spherical probe of 0.7 Å radius, focusing our analysis on the major branches of tunnels established in the dehalogenase family i.e. P1, P2, and P3,^45,68^ as well as recently explored side tunnel (ST).^68,69^ In addition to this, we also compared the bottleneck radii (BR), max BR and tunnel length to discern any significant differences in the structural properties of the tunnels identified by the two water models (**Figure 2 & Table S3A**).

**Figure 2:**
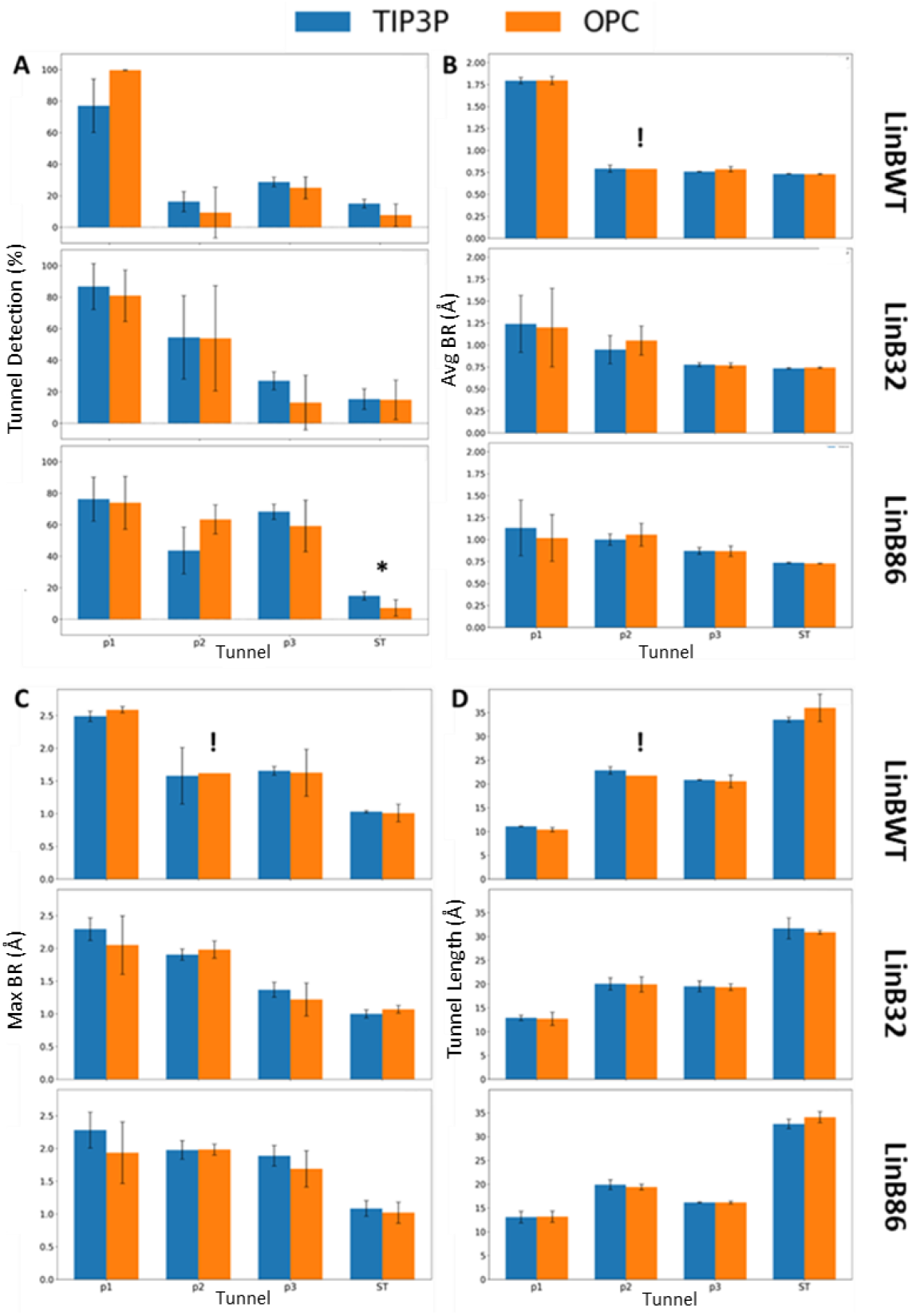
Frequency of tunnel detection and geometric characteristics for main and auxiliary tunnels for the three systems in TIP3P and OPC water models. **A**) Comparison of the frequency of tunnel detection (BR >0.7 Å) for the P1, P2, P3, and ST. Comparison of **B**) the average bottleneck radii, **C**) the maximum bottleneck radii, and **D**) the lengths of these four tunnels. Data represents average±standard deviation calculated from three or four MD simulations of LinBWT and its variants, respectively (see **Table S3A**). Statistically significant differences between OPC and TIP3P are indicated by asterisks above the bar plots. See also **Tables S3B-D** for details on the statistical test. The exclamation mark indicates the inability to conduct statistical tests since the tunnel was found in one replicate.

In LinBWT, the main tunnel (P1) could be detected for most of the simulation time, irrespective of the water model used **(Figure 2A)** with no statistically significant difference **(Table S3B)**. In LinB32 and LinB86 simulations, the effect of the L177W mutation hampering the opening of P1 tunnel was consistent between the water models, i.e., the average BR was reduced for P1 tunnel compared to LinBWT **(Figure 2B & Tables S3B-D)**. Also, the more frequent opening of the engineered P3 tunnel was consistently detected in LinB86 simulations with both water models **(Figure 2A & Table S3D)**, in line with the previous observations.^45^ Furthermore, the ST exhibited the lowest frequency of occurrence in all three systems with both water models **(Figure 2A)**. There was a statistically significant difference in the detection of this tunnel between simulations of LinB86 with OPC and TIP3P **(Table S3D**). Still, both models correctly ranked ST as the most rare. Finally, the average length of P1, P2, P3, and side tunnels was in excellent agreement between the water models **(Figure 2D & Table S3B-D)**. A detailed comparison of the tunnel detection and geometric characteristics (average BR, maximal BR and average length) of all tunnel clusters identified in OPC and TIP3P simulations can be found in **Tables S2A-C**. Overall, this data suggests that both water models can detect similar tunnel networks with only minor differences and provide comparable inferences about the effect of mutations on the tunnels.

### 3.2 Comparison of P1 bottleneck dimension between the TIP3P and OPC

The architecture of enzyme access tunnels plays a crucial role in determining ligand specificity, reaction kinetics, and overall enzyme stability.^70^ Narrower tunnel bottlenecks, while potentially limiting ligand transport, can reduce active site solvation and increase the probability of productive binding.^70^ The residues lining these tunnels also contribute by lowering initial entropy and facilitating effective interactions between the ligand and the active site.^70^ For instance, in the DhaA enzyme, a single point mutation (Y176A) that altered the bottleneck of the main tunnel led to a significant widening of the tunnel, which accelerated the binding of a fluorescent probe by three orders of magnitude.^71^ Due to the vital role of tunnel bottlenecks, we investigated the effect of water models on bottleneck geometry, focusing our detailed analysis on the main tunnel i.e. P1.

Data for tunnel geometries in LinBWT revealed that the main tunnel in both water models was qualitatively comparable with higher frequency observed for BR between 1.4 - 2.1 Å **(Figure 3)**. Similarly, the overall trends in the frequency of tunnel geometries in both water models were comparable in LinB32 simulations. The higher frequency distributions of BR were shifted to the lower end of the spectrum while a reasonable frequency was observed for 1.4 - 2.0 Å. For LinB86, similar profiles were seen for both water models with very high frequency distributions observed for shorter BR which is consistent with the closing of P1 tunnel for this variant. It is important to note that higher standard deviations could be observed for LinB32 & LinB86 which can be accounted for by including the two extremes, i.e. open and closed states. Although t-tests performed indicated no statistically significant difference for the water models in any of the BR ranges **(Table S4)**, one can observe a trend; simulations with OPC often resulted in narrower tunnels with 0.7 – 0.9 Å radius and less often produced open tunnels with radius ranging from 1.4- 2.0 Å **(Figure 3)**. In summary, these results indicate that the distribution of radii of the primary P1 tunnel was reasonably well maintained in simulations using either of the two water models.

**Figure 3:**
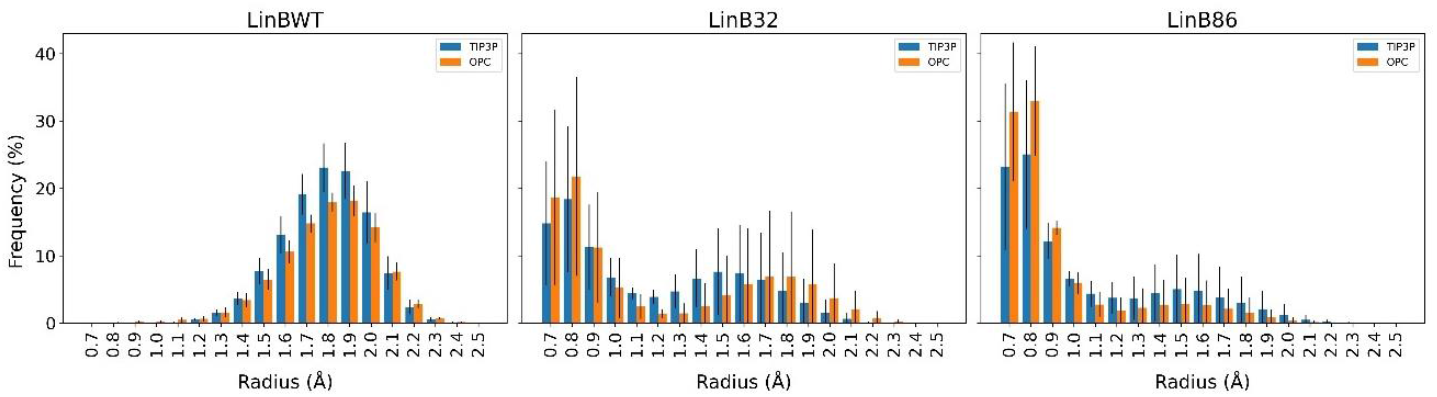
Bottleneck radii distribution of P1 tunnel found in the LinB variants simulated in TIP3P and OPC water models. Data represents average±standard deviation calculated from three or four MD simulations of LinBWT and its variants, respectively. Information on the statistical significance of the differences observed between the water models is available in **Table S4**.

### 3.3 Comparison of Duration of Tunnel Opening between the TIP3P and OPC

To understand the effect of water models on the stability of the functionally relevant open state of tunnels, we monitored the continuous duration of the main tunnel open states (≥ 1.4 Å) before they close again (< 1.4 Å) **(Figure 4)**. Although a recent study has identified a 0.7 Å radius as functionally relevant for tunnels of α/β-hydrolase superfamily,^72^ we opted for a 1.4 Å threshold to assess how tunnel openness might influence ligand interactions specifically. This threshold allows us to capture a broader range of tunnel dynamics, which is crucial for understanding the potential impact on ligand binding and transport. Unsurprisingly, opening events in the P1 tunnel were predominantly concluded in timescales between 20-100 ps in all simulated systems. However, we sporadically observed openings with a duration between 1-8 ns. Tunnel opening data for LinBWT revealed that the opening stability of both main tunnels was similar in both water models. A statistically significant difference could be observed for shorter-duration tunnel openings with these events being more frequent in OPC than TIP3P **(Figure 4A & Table S5)**.

**Figure 4:**
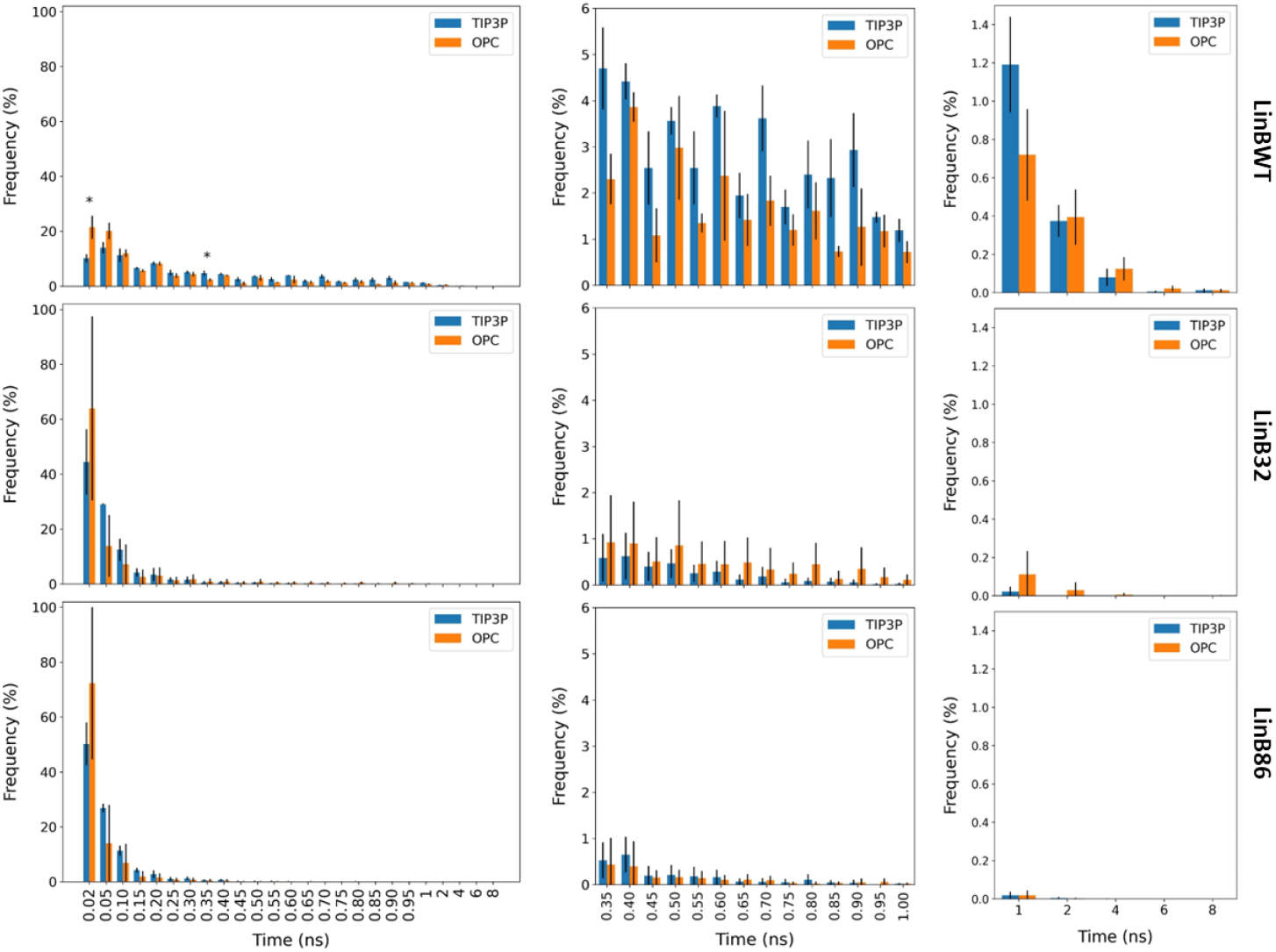
Duration of open states (BR ≥1.4 Å) of main P1 tunnel found in studied conformations of LinB variants simulated in TIP3P and OPC water models. **A**) The duration of open states spanning 0.02-8 ns. Data represents average±standard deviation calculated from three or four MD simulations of LinBWT and its variants, respectively. Statistically significant differences between OPC and TIP3P are indicated by asterisks above the bar plots. See also **Table S5** for details on the statistical test. **B**) A closer look at the duration of open states within the 0.35-8 ns range, while **C**) highlights the 1-8 ns timescale.

While for slightly higher duration opening events were more frequent in TIP3P than OPC. In LinB32, we observed a marked increase in the duration of tunnel openings in OPC water models, reaching up to 6 ns in OPC while being ∼ 1.5 ns for TIP3P. This observation is supported by the overestimated permeability of TIP3P water molecules seen in aquaporin AQP1^39^ and the underestimated binding affinity of these molecules in the interior of the Concanavalin A protein,^38^ which could diminish their contribution to stabilizing the open state. Similar to the situation in LinBWT, the differences in the opening duration in LinB86 were comparable between the two models, including the stabilities of the tunnel with the longest opening period of 1.5-2.0 ns. To sum up, the water models can markedly affect the stabilization of the open tunnel states, which can have severe implications for the effectiveness of their ability to transport small molecules. However, this observation is system-specific. Notably, the analysis indicates a preference for long-tailed events in OPC compared to TIP3P for LinB32. Conversely, TIP3P exhibited a higher frequency of long-tailed events for LinBWT and LinB86, although these differences were not statistically significant. This suggests that the choice of water model should be made with careful consideration of the specific system under study. A consistent trend observed in all the simulated systems was the prevalence of extremely short-duration events in OPC than in TIP3P. When evaluating trends in the three systems, the most distinct difference could be observed for the long-tailed events, wherein the frequency as well as the duration of the prolonged opened state was much more in comparison to LinB32 & LinB86, approximately 5 and 7 ns in TIP3P and OPC, respectively **(Figure 4C)**.

### 3.4 Assessment of Tunnel Cluster Sampling and Detection Rates With the TIP3P and OPC

To understand the effect of water models on the detection rates and sampling of tunnel clusters, we analyzed the performance of TIP3P and OPC across the three systems. In general, simulations using TIP3P detected more tunnel clusters and did so earlier **(Figure 5)**. The difference was the most pronounced in LinBWT compared to LinB32 and LinB86. This indicates that the TIP3P water model is more efficient in rapidly identifying tunnel clusters, potentially attributable to its dynamic properties. This can also be evidenced in **Table S1**, where the molecular gate governing P1 tunnel geometry is consistently more likely open in TIP3P compared to OPC for LinB32 and LinB86. Although TIP3P simulations led to the detection of a higher number of clusters within each system, these clusters tended to be the same set repeatedly. This suggests that TIP3P has a propensity for quickly identifying a consistent set of clusters. On the other hand, OPC demonstrated a distinct advantage by detecting more unique tunnel clusters, across all studied systems.

**Figure 5:**
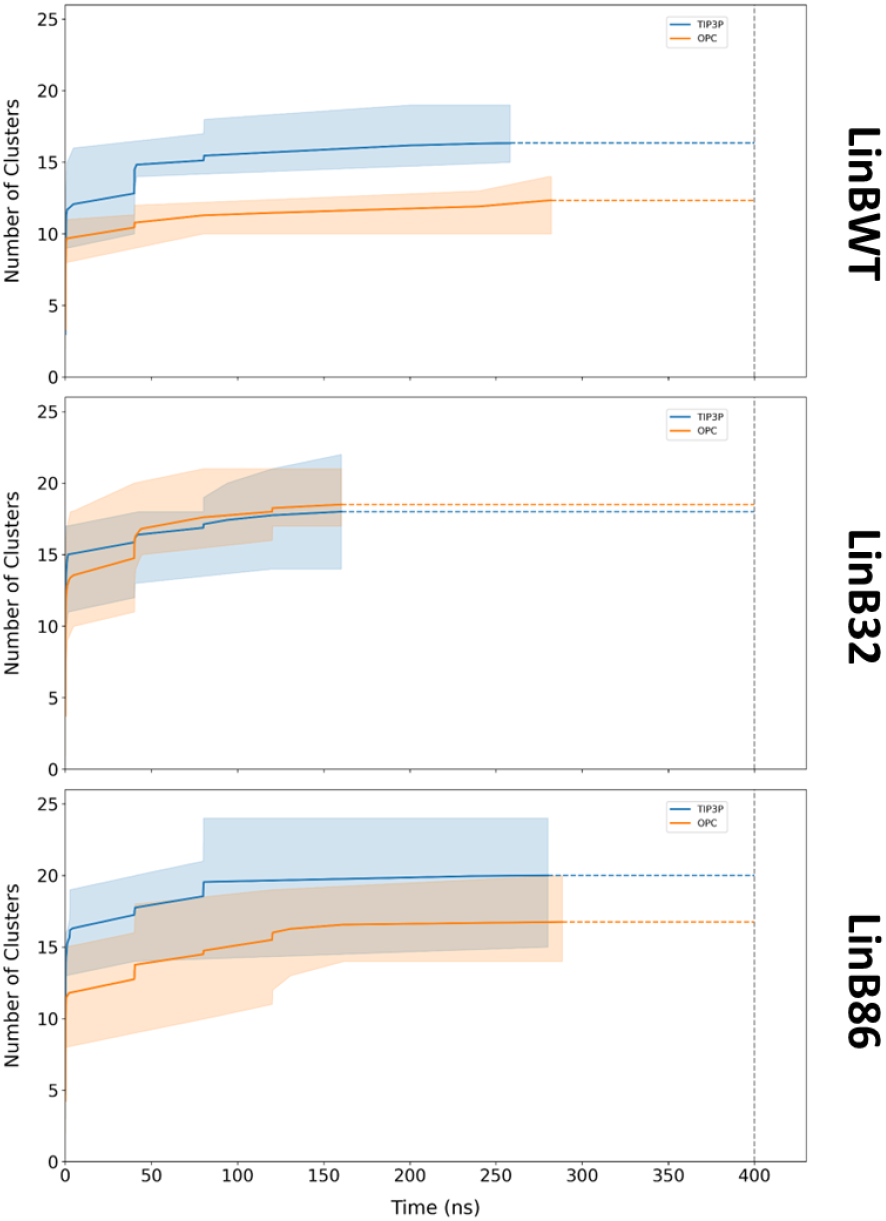
Average time taken for identification of tunnel clusters in the studied conformations of LinB variants simulated in TIP3P and OPC water models. The solid line represents the average number of clusters identified across three or four MD simulations of LinBWT and its variants, respectively. The shaded region represents the range of the number of clusters identified across all simulations. The vertical dotted line denotes the culmination of the simulation. The start of the horizontal dotted line (blue and orange) highlights the time after which no new cluster was identified in any of the simulation replicas.

This diversity in detection suggests that OPC provides a more comprehensive sampling of the tunnel network within the enzyme systems. Overall, OPC identified a total of 35 distinct tunnel clusters, while TIP3P detected 32 clusters. This marginal difference highlights that while TIP3P is efficient in terms of speed and frequency, OPC excels in providing a more thorough exploration albeit comparatively slowly. Notably, the unique clusters identified by OPC possessed tunnel cluster IDs (CIDs) 26, 27, and 36. CID 26 was identified in one simulation of LinBWT and two of LinB86; CID 27 emerged in one simulation each of LinBWT and LinB86, while CID 36 was detected in two simulations of LinB32. This distribution indicates that OPC’s ability to uncover these clusters extends across all three systems and is not merely an artifact of a particular system. Upon closer examination, it was found that all three CIDs corresponded to branches of the main tunnels. CID 26 appeared in approximately 2% of the frames, CID 27 in about 1.5%, and CID 36 in roughly 0.9%. Despite the relatively low frequency of occurrence, these superclusters represent unique extensions of the main tunnel and may be particularly valuable in longer exploratory simulations, where the likelihood of observing less frequent but potentially important features is enhanced. This detection of unique clusters by OPC could be crucial for understanding the full range of tunnel dynamics and their potential functional implications. While TIP3P and OPC each have their strengths, the data indicates that the choice of water model should be tailored to the specific objectives of the research.

## 4. CONCLUSIONS

Herein, we performed a series of MD simulations of LinBWT and its two engineered variants in TIP3P and OPC water models. Our simulations showed that structural ensembles of these proteins behaved in similar manners, with both water models exploring almost identical conformations and resulting in the detection of similar networks of transport tunnels. With these proteins, the OPC model simulations exhibited increased success in identifying narrower tunnels over TIP3P simulations. However, we also observed marked sensitivity of the stability of open states to the water model used, making their choice important when ligand binding and unbinding kinetics are being investigated, e.g., when estimating residence times of inhibitors within the drug discovery frameworks.^73–75^ Given the overestimated diffusivity of bulk TIP3P water molecules and its unrealistic migration rates via aquaporin AQP1, we believe the OPC water model represents a better choice under these circumstances. With this in mind, future research will expand to analyze explicitly the transport of water molecules via enzyme tunnels using different models with tools like AQUA-DUCT.^76^ Finally, we would like to highlight the 3-point TIP3P model is an equally suitable choice as OPC for the investigation of tunnel network geometry in scenarios where available computational resources or incompatibility with a protein force field may prohibit the use of the more costly 4-point OPC model.

## CRediT Authorship Contribution Statement

**A.S**.: Conceptualization, Data curation, Formal analysis, Investigation, Methodology, Software, Validation, Visualization, Writing – original draft, Writing – review & editing; **N.A**.: Conceptualization, Data curation, Formal analysis, Investigation, Methodology, Software, Writing–original draft; **J.B**.: Conceptualization, Data curation, Funding acquisition, Methodology, Project administration, Resources, Software, Supervision, Writing – original draft, Writing – review & editing.

## Acknowledgments

This work was supported by the National Science Centre, Poland (grant no. 2017/26/E/NZ1/00548). N.A. acknowledge the ANM postdoc grant (no. ANM_OSI_PG_65) for financial support. The computations were performed at the Poznan Supercomputing and Networking Center. The authors are grateful to Bartłomiej Surpeta, Carlos Sequeiros-Borja, and Nishita Mandal, Adam Mickiewicz University, Poznan, Poland for critical reading of the manuscript and helpful discussions.

## Data Availability Statement

The data underlying this study are available in the published article, its supplementary information, and in the Zenodo repository at https://doi.org/10.5281/zenodo.7929605. The data were deposited as plain text, PDB formatted structural data and AMBER-formatted MD trajectories, which can be processed via numerous freely available SW packages, no tools with restricted access are needed.

## Abbreviations

BR: bottleneck radii
CID: Cluster ID
HLD: haloalkane dehalogenase
MD: molecular dynamics
OPC: optimal point charge
RDF: radial distribution function
RMSD: root-mean-square deviation
RMSF: residue-wise root mean square fluctuation
SPC: single point charge
ST: Side Tunnel
TIP: transferable intermolecular potential.

