## Supplementary Materials for "Impact of water models on structure and dynamics of enzyme tunnels"

***Text S1. The conformational state of cap domain gates in dehalogenases are structurally comparable between OPC and TIP3P.***

In a previous study using both experimental and simulation techniques, Brezovsky *et al.* discovered that LinB32 and LinB86 can adopt open and closed conformations due to the marked movement of one of the cap domain helices after the hydrogen bond between the Asp147 and mutated Trp177 is broken (Brezovsky et al., 2016; Kokkonen et al., 2018). Moreover, these open and closed states were shown to be associated with very different enzymatic efficiency (Kokkonen et al., 2018). To identify the open and closed states of these proteins in our simulations, we calculated the percentage of time that the gate between residues 147 and 177 remained open for water molecules in each system (**Table S1**). Due to the lack of hydrogen bonds stabilizing the closed gate conformation, each of the three initial simulations of LinBWT with both water models exhibited an almost exclusively open state. In the case of the mutants, additional simulations had to be run to obtain at least two simulations that sampled the opened states for at least 20 % of the time. Furthermore, we found that to discover open states for LinB32 with the OPC water model required approximately twice as many screening simulations in comparison to other systems. This is consistent with a slower sampling of the opening process in OPC and the previously established enhanced structural stability of the cap-domain helical gate in this mutant.(Brezovsky et al., 2016; Kokkonen et al., 2018) The final set of two simulations with the most often open state (LinB32-Open and LinB86-Open) and the two most often closed simulations (LinB32-Closed and LinB86-Closed) were assembled for each water model (**Table S1**).

**Table S1. Opening of molecular gate between cap domain helices for water molecules.**

| Simulation Replica # | Percent of simulations with minimum distance between ASP147 and TRP177 > 4 Å | | | | | |
|---|---|---|---|---|---|---|
| System | LinBWT | | LinB32 | | LinB86 | |
| | TIP3P | OPC | TIP3P | OPC* | TIP3P | OPC |
| 1 | 95 | 99 | 12 | 1 | 23 | 5 |
| 2 | 97 | 97 | 24 | 61 | 4 | 6 |
| 3 | 98 | 98 | 24 | 2 | 32 | 4 |
| 4 | | | 24 | 18 | 23 | 11 |
| 5 | | | 43 | 1 | 31 | 21 |
| 6 | | | 5 | 6 | 38 | 28 |
| 7 | | | 6 | 2 | 35 | |
| 8 | | | 28 | 1 | 3 | |
| 9 | | | 20 | 1 | 41 | |
| 10 | | | | 1 | | |
| 11 | | | | 1 | | |
| 12 | | | | 25 | | |
| 13 | | | | 2 | | |
| 14 | | | | 1 | | |
| 15 | | | | 1 | | |
| 16 | | | | 1 | | |
| 17 | | | | 9 | | |
| 18 | | | | 3 | | |

The simulations chosen to represent open and closed states for further investigation are highlighted in yellow and red, respectively. The simulations were performed in batches of three replicates, until the two suitable representatives were found, with the exception of LinB32-OPC, in which after three such batches (3x3 simulations), additional 9 simulations were executed. * In case of LinB32-OPC, the simulations representing the closed state were selected from among the most closed ones, by also minimizing the deviation in the investigated distances.

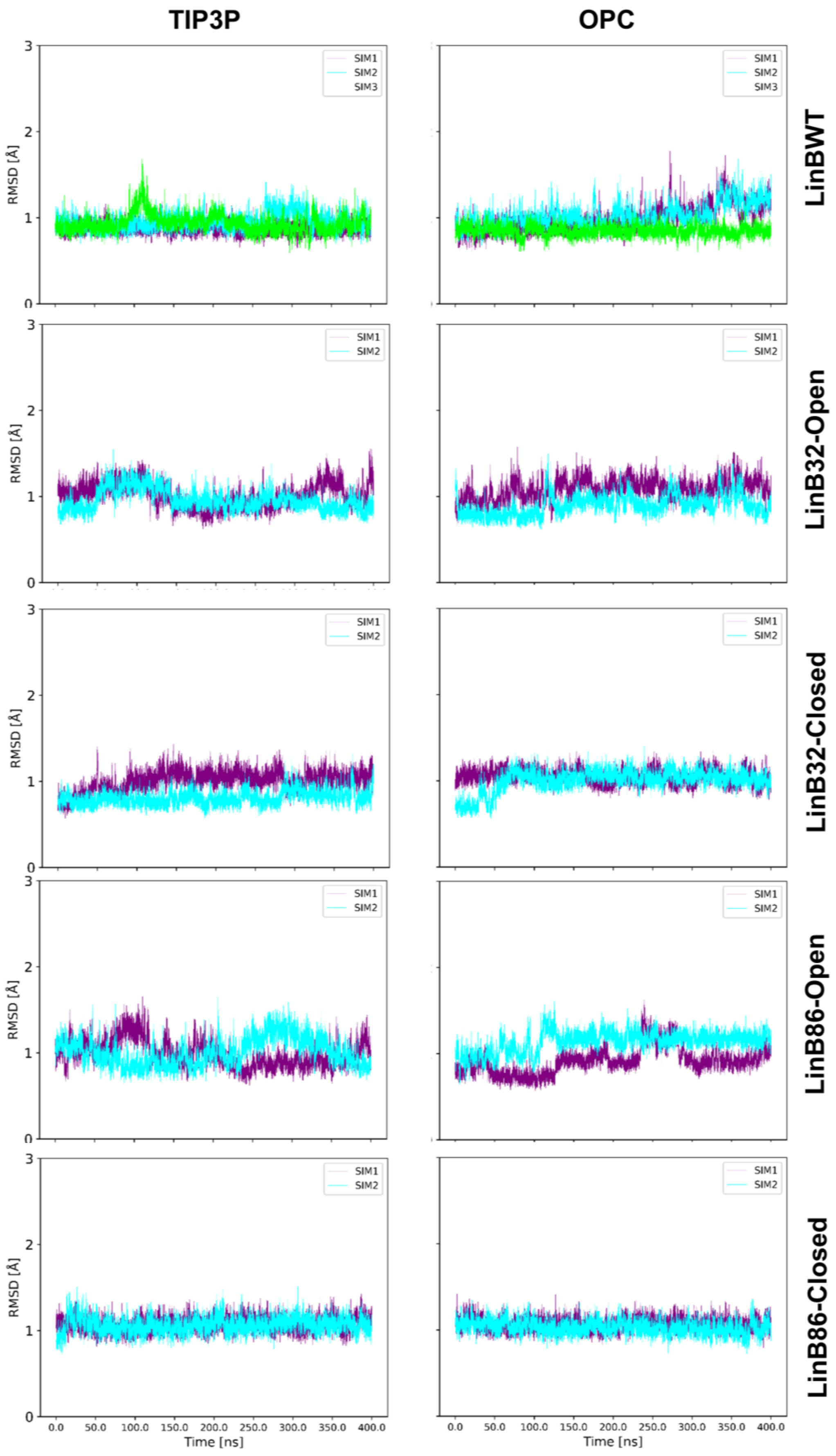


**Figure S1. Time evolution of backbone RMSD in production phase investigated simulations in TIP3P and OPC water models.**

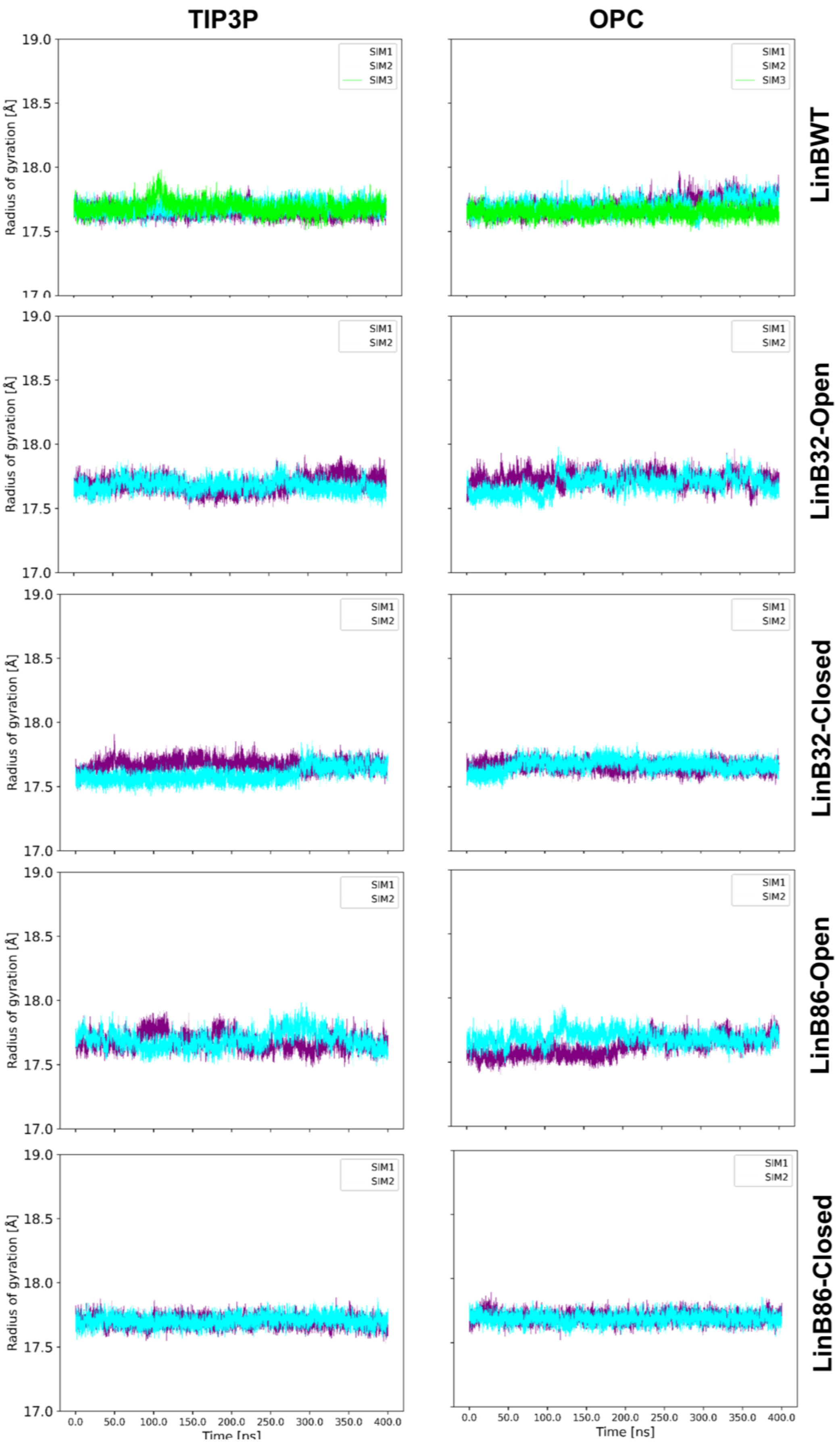


**Figure S2. Time evolution of backbone radius of gyration in production phase of investigated simulations in TIP3P and OPC water models.**

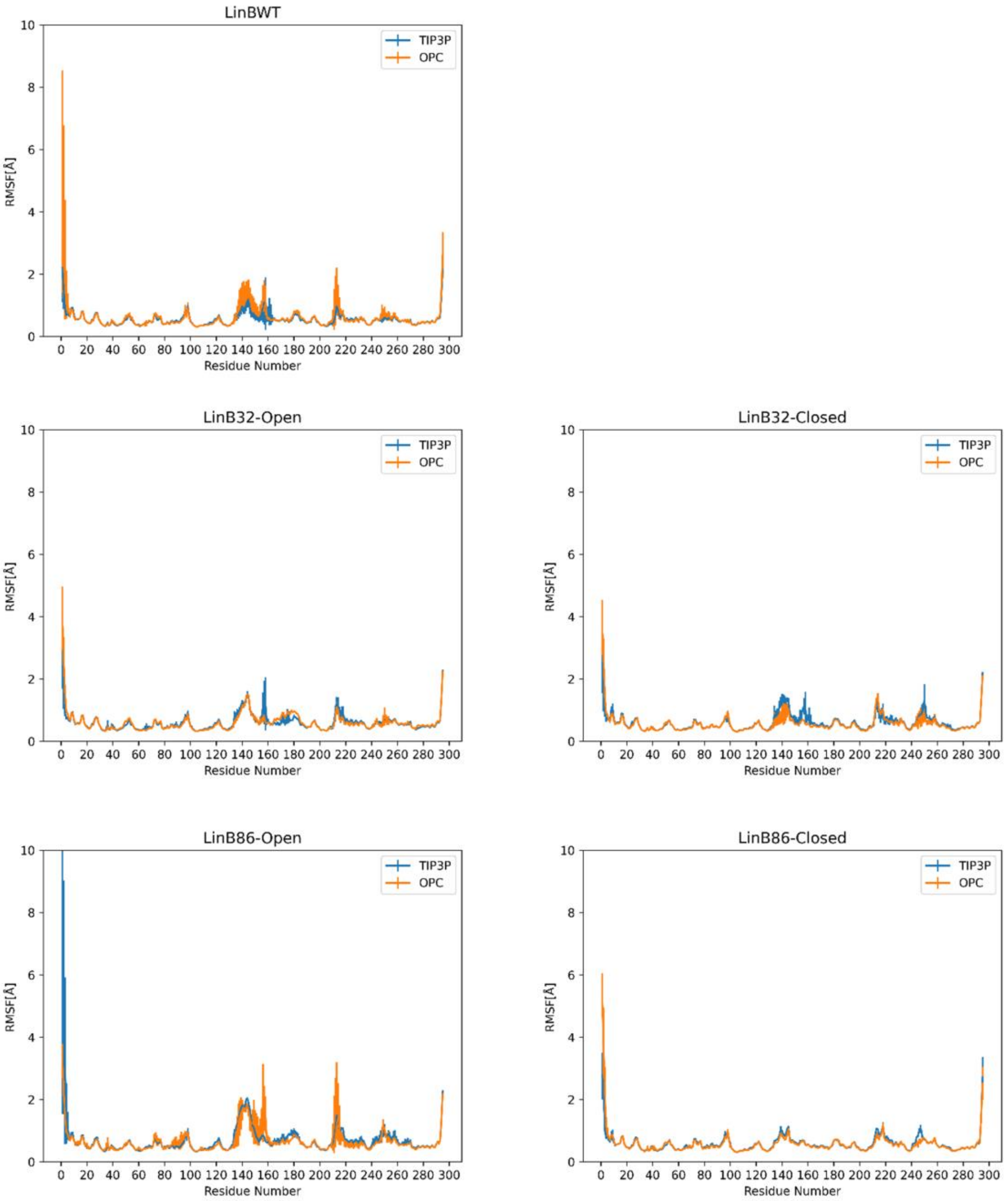


**Figure S3. Average backbone RMSF of residues in production phase investigated simulations in TIP3P and OPC water models.**

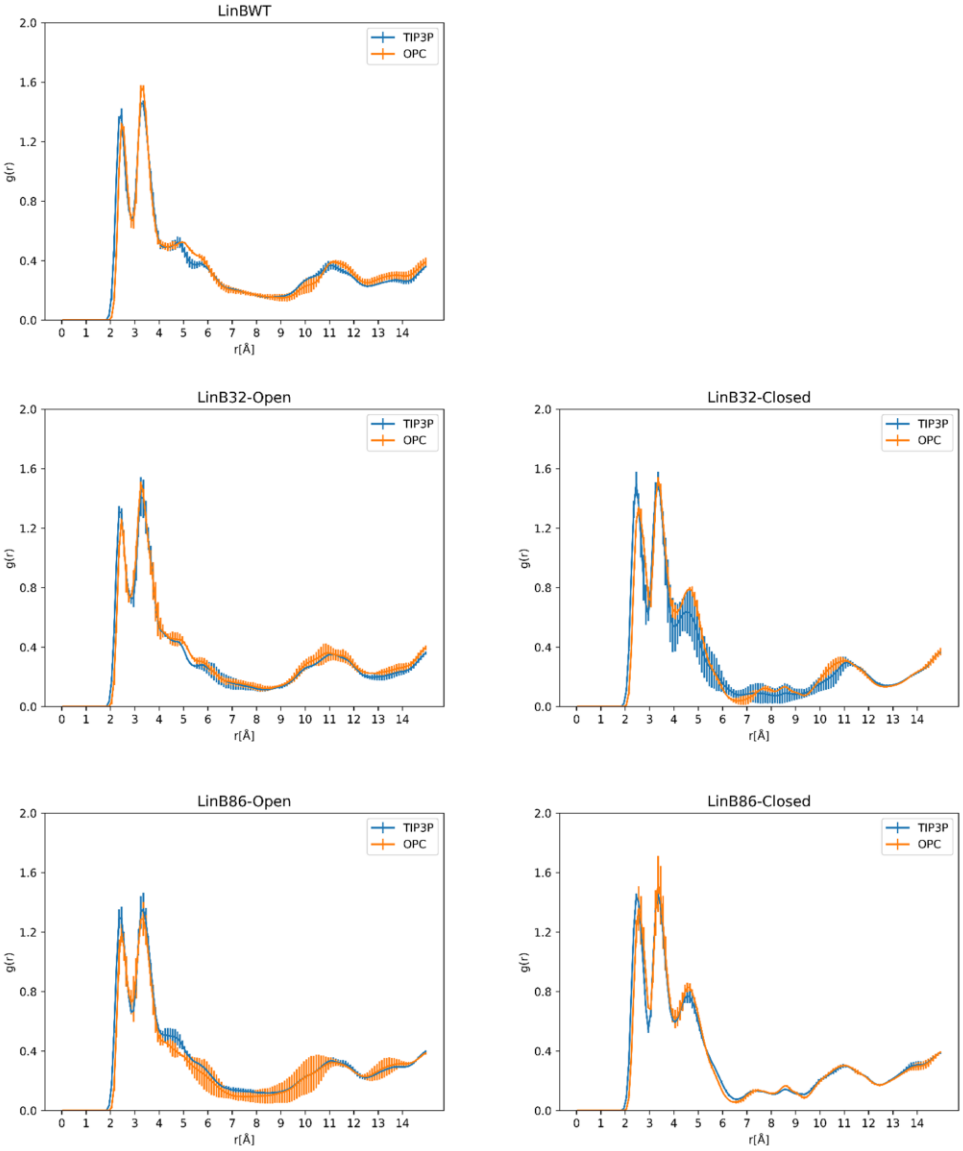


**Figure S4. Average radial distribution function of water molecules around CG atom of Asp108 in investigated simulations in TIP3P and OPC water models.**

**Table S2A. Comparison of characteristcis of all tunnel clusters identified in TIP3P and OPC water model in LinBWT.**

| **Cluster** | **No. of Simulations** | | **Total Frames** | | **Average Bottleneck Radii (Å)** | | **Maximum Bottleneck Radii (Å)** | | **Average Length (Å)** | |
|---|---|---|---|---|---|---|---|---|---|---|
| CID | OPC | TIP3P | OPC | TIP3P | OPC | TIP3P | OPC | TIP3P | OPC | TIP3P |
| *1* | 1 | 1 | 59787 | 46232 | 1.797 ± 0.224 | 1.796 ± 0.199 | 2.642 | 2.552 | 10.401 ± 1.655 | 11.079 ± 1.132 |
| *2* | 2 | 2 | 8080 | 12805 | 0.818 ± 0.150 | 1.791 ± 0.194 | 2.17 | 2.378 | 17.759 ± 3.439 | 11.368 ± 1.135 |
| *3* | 1 | 3 | 5568 | 9758 | 0.791 ± 0.091 | 0.803 ± 0.128 | 1.62 | 1.839 | 21.780 ± 2.305 | 22.644 ± 2.442 |
| *4* | 3 | 3 | 15018 | 17240 | 0.787 ± 0.107 | 0.759 ± 0.063 | 1.924 | 1.712 | 20.668 ± 3.497 | 20.864 ± 2.503 |
| *5* | 3 | 3 | 7423 | 14224 | 0.766 ± 0.072 | 0.775 ± 0.076 | 1.341 | 1.361 | 23.755 ± 3.175 | 23.875 ± 3.429 |
| *6* | 3 | 3 | 21329 | 26140 | 0.758 ± 0.054 | 0.763 ± 0.058 | 1.16 | 1.205 | 26.266 ± 3.700 | 26.075 ± 3.362 |
| *7* | 3 | 2 | 24739 | 5504 | 0.842 ± 0.147 | 0.800 ± 0.106 | 1.823 | 1.676 | 19.125 ± 2.818 | 19.177 ± 2.738 |
| *8* | 2 | 3 | 6339 | 11458 | 0.962 ± 0.252 | 0.828 ± 0.143 | 2.173 | 2.046 | 21.649 ± 2.576 | 20.395 ± 2.603 |
| *9* | - | - | - | - | - | - | - | - | - | - |
| *10* | - | - | - | - | - | - | - | - | - | - |
| *11* | 2 | 3 | 4577 | 8991 | 0.731 ± 0.037 | 0.734 ± 0.037 | 1.105 | 1.043 | 35.528 ± 7.719 | 33.457 ± 6.943 |
| *12* | 3 | 3 | 4405 | 5835 | 0.730 ± 0.034 | 0.731 ± 0.034 | 1.365 | 1.081 | 30.687 ± 7.595 | 29.282 ± 6.503 |
| *13* | - | 1 | - | 1660 | - | 0.740 ± 0.042 | - | 0.947 | - | 32.983 ± 4.853 |
| *14* | - | 1 | - | 1128 | - | 0.759 ± 0.061 | - | 1.112 | - | 27.009 ± 3.300 |
| *15* | 1 | 2 | 3963 | 2770 | 0.799 ± 0.126 | 0.778 ± 0.097 | 1.797 | 1.692 | 18.534 ± 3.092 | 19.225 ± 3.178 |
| *16* | 1 | 1 | 1539 | 6788 | 0.803 ± 0.096 | 0.782 ± 0.088 | 1.26 | 1.265 | 31.352 ± 4.510 | 31.889 ± 4.992 |
| *17* | - | 3 | - | 5224 | - | 0.739 ± 0.038 | - | 1.04 | - | 31.667 ± 5.387 |
| *18* | 2 | 2 | 2681 | 3037 | 0.729 ± 0.030 | 0.729 ± 0.029 | 0.914 | 0.946 | 30.106 ± 3.796 | 31.131 ± 3.779 |
| *19* | - | 3 | - | 5881 | - | 0.740 ± 0.042 | - | 1.222 | 36.091 ± 5.925 | - |
| *20* | - | 1 | - | 1395 | - | 0.754 ± 0.052 | - | 1.12 | -- | 21.029 ± 2.350 |
| *21* | 3 | - | 8527 | - | 0.735 ± 0.037 | - | 1.105 | - | 33.223 ± 6.178 | - |
| *22* | - | 2 | - | 2987 | - | 0.731 ± 0.031 | - | 0.946 | - | 48.677 ± 5.075 |
| *23* | - | - | - | - | - | - | - | - | - | - |
| *24* | - | 2 | - | 2608 | - | 0.723 ± 0.023 | - | 0.897 | - | 36.440 ± 6.085 |
| *25* | - | - | - | - | - | - | - | - | - | - |
| *26* | - | - | - | - | - | - | - | - | - | - |
| *27* | - | - | - | - | - | - | - | - | - | - |
| *28* | - | - | - | - | - | - | - | - | - | - |
| *29* | - | 2 | - | 2873 | - | 0.728 ± 0.029 | - | 0.951 | - | 47.479 ± 6.664 |
| *30* | - | 1 | - | 1095 | - | 0.725 ± 0.025 | - | 0.856 | - | 64.230 ± 5.994 |
| *31* | 1 | - | 1678 | - | 0.752 ± 0.062 | - | 1.258 | | 25.822 ± 4.101 | - |
| *32* | - | - | - | - | - | - | - | - | - | - |
| *33* | - | - | - | - | - | - | - | - | - | - |
| *34* | - | - | - | - | - | - | - | - | - | - |
| *35* | 1 | - | 1066 | - | 0.726 ± 0.025 | - | 0.902 | - | 36.695 ± 5.371 | - |
| *36* | - | - | - | - | - | - | - | - | - | - |

**Table S2B. Comparison of characteristics of all tunnel clusters identified in TIP3P and OPC water model in LinB32.**

| **Cluster** | **No. of Simulations** | | **Total Frames** | | **Average Bottleneck Radii (Å)** | | **Maximum Bottleneck Radii (Å)** | | **Average Length (Å)** | |
|---|---|---|---|---|---|---|---|---|---|---|
| CID | OPC | TIP3P | OPC | TIP3P | OPC | TIP3P | OPC | TIP3P | OPC | TIP3P |
| *1* | 4 | 4 | 64769 | 69368 | 1.260 ± 0.461 | 1.276 ± 0.398 | 2.482 | 2.448 | 12.514 ± 2.465 | 12.849 ± 1.995 |
| *2* | 4 | 4 | 52566 | 60032 | 1.261 ± 0.436 | 1.239 ± 0.375 | 2.42 | 2.558 | 13.950 ± 2.789 | 14.052 ± 2.627 |
| *3* | 4 | 4 | 43122 | 43578 | 1.111 ± 0.272 | 1.005 ± 0.273 | 2.108 | 1.996 | 19.226 ± 2.153 | 19.670 ± 2.873 |
| *4* | 3 | 4 | 10481 | 21558 | 0.787 ± 0.090 | 0.774 ± 0.081 | 1.503 | 1.533 | 18.990 ± 3.478 | 19.640 ± 3.127 |
| *5* | 4 | 4 | 49873 | 34884 | 0.860 ± 0.118 | 0.849 ± 0.134 | 1.555 | 1.763 | 22.296 ± 2.709 | 22.391 ± 3.022 |
| *6* | 4 | 4 | 46562 | 35277 | 0.788 ± 0.077 | 0.779 ± 0.073 | 1.284 | 1.296 | 26.210 ± 3.910 | 25.880 ± 4.041 |
| *7* | 3 | 3 | 17055 | 26932 | 0.813 ± 0.119 | 0.869 ± 0.147 | 1.736 | 1.82 | 16.749 ± 2.462 | 16.667 ± 2.401 |
| *8* | 2 | - | 17240 | - | 1.041 ± 0.287 | - | 2.251 | - | 17.812 ± 3.611 | - |
| *9* | 2 | 1 | 27017 | 15124 | 0.857 ± 0.136 | 0.860 ± 0.130 | 1.559 | 1.567 | 21.816 ± 2.866 | 23.945 ± 3.467 |
| *10* | 3 | 1 | 16810 | 7826 | 0.761 ± 0.062 | 0.770 ± 0.071 | 1.288 | 1.635 | 27.598 ± 5.072 | 27.457 ± 4.806 |
| *11* | 3 | 4 | 11846 | 12267 | 0.745 ± 0.048 | 0.737 ± 0.041 | 1.113 | 1.04 | 30.965 ± 6.797 | 31.142 ± 7.088 |
| *12* | 2 | 4 | 5620 | 8065 | 0.745 ± 0.048 | 0.734 ± 0.038 | 1.163 | 1.09 | 27.842 ± 5.201 | 28.278 ± 6.424 |
| *13* | 2 | 3 | 11657 | 8002 | 0.842 ± 0.118 | 0.792 ± 0.097 | 1.537 | 1.471 | 28.050 ± 3.009 | 30.537 ± 5.156 |
| *14* | 2 | 1 | 5734 | 4920 | 0.770 ± 0.068 | 0.772 ± 0.069 | 1.141 | 1.239 | 29.250 ± 5.037 | 28.495 ± 4.726 |
| *15* | 2 | 1 | 2956 | 1975 | 0.785 ± 0.106 | 0.779 ± 0.093 | 1.885 | 1.758 | 18.024 ± 3.286 | 18.980 ± 3.827 |
| *16* | 2 | 3 | 8428 | 10811 | 0.791 ± 0.091 | 0.785 ± 0.088 | 1.574 | 1.337 | 30.635 ± 4.591 | 30.957 ± 5.487 |
| *17* | 3 | 3 | 7067 | 4653 | 0.740 ± 0.039 | 0.740 ± 0.039 | 1.007 | 0.984 | 31.539 ± 5.761 | 30.839 ± 5.511 |
| *18* | 2 | 3 | 4032 | 4785 | 0.731 ± 0.031 | 0.729 ± 0.030 | 0.923 | 0.939 | 30.871 ± 4.463 | 30.214 ± 3.781 |
| *19* | 3 | 2 | 4216 | 5376 | 0.744 ± 0.046 | 0.744 ± 0.045 | 1.102 | 1.143 | 35.462 ± 6.302 | 36.517 ± 5.419 |
| *20* | 3 | 2 | 8950 | 2533 | 0.751 ± 0.059 | 0.750 ± 0.051 | 1.66 | 1.132 | 20.571 ± 2.916 | 20.284 ± 2.677 |
| *21* | - | 1 | - | 1989 | - | 0.736 ± 0.035 | - | 0.967 | - | 31.833 ± 5.272 |
| *22* | 3 | 3 | 4913 | 5316 | 0.727 ± 0.029 | 0.731 ± 0.033 | 0.943 | 0.96 | 45.349 ± 5.752 | 46.348 ± 5.688 |
| *23* | 2 | - | 7083 | - | 0.830 ± 0.139 | - | 2.013 | - | 20.425 ± 5.095 | - |
| *24* | 2 | 2 | 5108 | 2868 | 0.730 ± 0.030 | 0.724 ± 0.025 | 0.935 | 0.918 | 33.762 ± 5.197 | 33.982 ± 5.871 |
| *25* | 1 | 1 | 1353 | 1123 | 0.745 ± 0.040 | 0.741 ± 0.037 | 0.94 | 0.921 | 36.151 ± 4.611 | 35.396 ± 4.475 |
| *26* | - | - | - | - | - | - | - | - | - | - |
| *27* | - | - | - | - | - | - | - | - | - | - |
| *28* | 2 | 1 | 2655 | 1347 | 0.731 ± 0.033 | 0.732 ± 0.033 | 0.997 | 0.983 | 34.946 ± 5.789 | 33.453 ± 4.995 |
| *29* | 2 | 2 | 3453 | 2219 | 0.732 ± 0.031 | 0.728 ± 0.029 | 0.901 | 0.912 | 46.285 ± 5.978 | 47.132 ± 6.949 |
| *30* | 1 | 2 | 1083 | 3423 | 0.723 ± 0.024 | 0.723 ± 0.023 | 0.921 | 0.893 | 55.470 ± 5.788 | 58.507 ± 7.628 |
| *31* | - | 1 | - | 1274 | - | 0.746 ± 0.053 | - | 1.142 | - | 24.360 ± 3.696 |
| *32* | 2 | 1 | 2748 | 1461 | 0.731 ± 0.031 | 0.730 ± 0.030 | 0.909 | 0.899 | 30.319 ± 5.121 | 29.555 ± 4.019 |
| *33* | - | 1 | - | 1865 | - | 0.738 ± 0.038 | - | 0.959 | - | 31.044 ± 5.375 |
| *34* | - | 2 | - | 2433 | - | 0.727 ± 0.025 | - | 0.872 | - | 35.695 ± 5.172 |
| *35* | 2 | - | 4029 | - | 0.740 ± 0.043 | - | 1.041 | - | 29.540 ± 6.371 | |
| *36* | - | - | - | - | - | - | - | - | - | - |

**Table S2C. Comparison of characteristics of all tunnel clusters identified in TIP3P and OPC water model in LinB86.**

| **Cluster** | **No. of Simulations** | | **Total Frames** | | **Average Bottleneck Radii (Å)** | | **Maximum Bottleneck Radii (Å)** | | **Average Length (Å)** | |
|---|---|---|---|---|---|---|---|---|---|---|
| **CID** | OPC | TIP3P | OPC | TIP3P | OPC | TIP3P | OPC | TIP3P | OPC | TIP3P |
| ***1*** | 4 | 4 | 59136 | 60970 | 1.060 ± 0.353 | 1.174 ± 0.413 | 2.4 | 2.566 | 13.014 ± 2.438 | 12.926 ± 2.251 |
| ***2*** | 4 | 4 | 48680 | 59330 | 0.904 ± 0.264 | 1.025 ± 0.335 | 2.292 | 2.393 | 15.158 ± 2.834 | 14.699 ± 2.933 |
| ***3*** | 4 | 4 | 50719 | 34867 | 1.067 ± 0.275 | 1.012 ± 0.247 | 2.074 | 2.114 | 19.411 ± 2.217 | 19.705 ± 2.046 |
| *4* | 4 | 4 | 47338 | 54551 | 0.862 ± 0.154 | 0.873 ± 0.163 | 1.938 | 2.09 | 16.118 ± 2.323 | 16.175 ± 2.208 |
| ***5*** | 4 | 4 | 44159 | 54282 | 0.880 ± 0.131 | 0.884 ± 0.133 | 1.569 | 1.655 | 23.610 ± 3.128 | 23.370 ± 3.536 |
| ***6*** | 4 | 4 | 33699 | 48859 | 0.794 ± 0.082 | 0.810 ± 0.092 | 1.239 | 1.334 | 28.636 ± 4.673 | 27.373 ± 4.539 |
| ***7*** | 4 | 3 | 13513 | 17568 | 0.873 ± 0.213 | 0.849 ± 0.140 | 1.937 | 1.78 | 16.661 ± 2.939 | 17.703 ± 2.842 |
| ***8*** | 3 | 4 | 9548 | 29945 | 0.886 ± 0.176 | 1.074 ± 0.304 | 2.03 | 2.34 | 17.986 ± 2.459 | 16.847 ± 3.086 |
| ***9*** | 2 | 3 | 29757 | 37342 | 0.907 ± 0.139 | 0.963 ± 0.168 | 1.629 | 1.997 | 21.823 ± 2.861 | 22.366 ± 2.966 |
| ***10*** | 2 | 4 | 22908 | 33024 | 0.778 ± 0.073 | 0.777 ± 0.075 | 1.254 | 1.296 | 27.789 ± 4.290 | 28.829 ± 5.231 |
| ***11*** | 3 | 4 | 5740 | 11888 | 0.729 ± 0.033 | 0.736 ± 0.042 | 1.204 | 1.26 | 34.029 ± 7.977 | 32.799 ± 7.653 |
| *12* | 2 | 4 | 2894 | 8777 | 0.731 ± 0.033 | 0.736 ± 0.042 | 0.984 | 1.211 | 29.478 ± 7.265 | 28.642 ± 6.322 |
| ***13*** | 2 | 1 | 11319 | 1179 | 0.980 ± 0.148 | 0.789 ± 0.094 | 1.579 | 1.239 | 23.260 ± 2.735 | 31.159 ± 5.691 |
| ***14*** | 2 | 4 | 4823 | 16227 | 0.756 ± 0.053 | 0.769 ± 0.069 | 1.095 | 1.27 | 32.092 ± 4.545 | 31.412 ± 5.912 |
| *15* | 2 | 1 | 3762 | 1105 | 0.808 ± 0.120 | 0.784 ± 0.094 | 1.502 | 1.702 | 16.653 ± 4.216 | 19.359 ± 3.596 |
| ***16*** | 1 | - | 2728 | - | 0.760 ± 0.066 | - | 1.54 | - | 32.231 ± 6.720 | - |
| ***17*** | 1 | 3 | 1465 | 6667 | 0.750 ± 0.046 | 0.741 ± 0.039 | 0.986 | 1.083 | 29.756 ± 6.165 | 30.744 ± 5.588 |
| ***18*** | 2 | 3 | 2940 | 4730 | 0.730 ± 0.031 | 0.733 ± 0.034 | 0.975 | 1.01 | 29.811 ± 4.105 | 30.019 ± 3.825 |
| ***19*** | 1 | 2 | 2737 | 3488 | 0.738 ± 0.040 | 0.745 ± 0.045 | 0.96 | 1.008 | 35.623 ± 4.562 | 36.571 ± 5.363 |
| ***20*** | - | - | - | - | - | - | - | - | - | - |
| ***21*** | 2 | 1 | 2674 | 2508 | 0.740 ± 0.041 | 0.742 ± 0.040 | 0.996 | 0.951 | 30.315 ± 5.578 | 32.163 ± 5.813 |
| ***22*** | 3 | 2 | 4227 | 3178 | 0.731 ± 0.033 | 0.733 ± 0.034 | 0.971 | 0.936 | 44.443 ± 4.528 | 45.809 ± 4.943 |
| ***23*** | - | 1 | - | 1266 | - | 0.836 ± 0.141 | - | 1.929 | - | 19.531 ± 4.279 |
| ***24*** | - | 2 | - | 3366 | - | 0.727 ± 0.027 | - | 0.953 | - | 35.371 ± 5.690 |
| ***25*** | 2 | 3 | 2308 | 3581 | 0.739 ± 0.037 | 0.742 ± 0.040 | 0.978 | 1.003 | 36.630 ± 5.781 | 35.261 ± 4.638 |
| ***26*** | 2 | - | 7501 | - | 0.762 ± 0.064 | - | 1.223 | - | 25.013 ± 5.051 | - |
| ***27*** | 1 | - | 4659 | - | 0.806 ± 0.104 | - | 1.69 | - | 16.325 ± 1.598 | - |
| ***28*** | 1 | 3 | 1047 | 3759 | 0.730 ± 0.033 | 0.728 ± 0.030 | 0.969 | 0.953 | 33.766 ± 5.353 | 36.014 ± 6.423 |
| ***29*** | 1 | 1 | 1101 | 1761 | 0.740 ± 0.040 | 0.732 ± 0.032 | 0.953 | 0.927 | 43.133 ± 5.116 | 45.637 ± 5.621 |
| ***30*** | 2 | 4 | 2898 | 5600 | 0.723 ± 0.024 | 0.724 ± 0.026 | 0.91 | 0.93 | 54.682 ± 5.257 | 58.203 ± 7.175 |
| ***31*** | 1 | - | 1391 | - | 0.765 ± 0.071 | - | 1.158 | - | 21.141 ± 3.602 | - |
| ***32*** | - | 1 | - | 1000 | - | 0.729 ± 0.028 | - | 0.929 | - | 31.945 ± 3.987 |
| ***33*** | 1 | 1 | 3052 | 1064 | 0.737 ± 0.039 | 0.747 ± 0.051 | 1.034 | 1.193 | 30.330 ± 6.032 | 31.877 ± 5.992 |
| *34* | - | - | - | - | - | - | - | - | - | - |
| ***35*** | - | 1 | - | 1010 | - | 0.727 ± 0.027 | - | 0.863 | - | 34.756 ± 5.796 |
| ***36*** | - | - | - | - | - | - | - | - | - | - |

**Table S3A. Mean and standard deviation (SD) values for the Tunnel Detection - Absolute Value (Frames), Average Bottleneck Radius (Avg_BR), Max Bottleneck Radius (Max_BR) and Length (Len) for LinB systems and the two water models.**

| *Tunnel* | *Frames* | *SD* | *Avg_BR* (Å) | *SD* | *Max_BR* (Å) | *SD* | *Len* (Å) | *SD* | *Model* | *System* |
|---|---|---|---|---|---|---|---|---|---|---|
| *p1* | 19929.000 | 24.880 | 1.797 | 0.045 | 2.588 | 0.048 | 10.401 | 0.440 | OPC | LinBWT |
| *p2* | 1856.000 | 3214.686 | 0.791 | - | 1.62 | - | 21.780 | - | OPC | LinBWT |
| *p3* | 5006.000 | 1369.202 | 0.787 | 0.029 | 1.627 | 0.358 | 20.561 | 1.300 | OPC | LinBWT |
| *ST* | 1525.667 | 1422.303 | 0.730 | 0.006 | 1.010 | 0.134 | 35.998 | 2.889 | OPC | LinBWT |
| *p1* | 15410.667 | 3384.586 | 1.794 | 0.037 | 2.489 | 0.079 | 11.091 | 0.089 | TIP3P | LinBWT |
| *p2* | 3252.667 | 1269.896 | 0.792 | 0.040 | 1.579 | 0.432 | 22.845 | 0.790 | TIP3P | LinBWT |
| *p3* | 5746.667 | 612.670 | 0.759 | 0.003 | 1.656 | 0.065 | 20.865 | 0.080 | TIP3P | LinBWT |
| *ST* | 2997.000 | 544.025 | 0.734 | 0.003 | 1.030 | 0.019 | 33.516 | 0.506 | TIP3P | LinBWT |
| *p1* | 16192.250 | 3263.929 | 1.198 | 0.447 | 2.051 | 0.446 | 12.710 | 1.376 | OPC | LinB32 |
| *p2* | 10780.500 | 6661.317 | 1.051 | 0.165 | 1.983 | 0.132 | 19.948 | 1.586 | OPC | LinB32 |
| *p3* | 2620.250 | 3464.777 | 0.768 | 0.026 | 1.219 | 0.250 | 19.356 | 0.708 | OPC | LinB32 |
| *ST* | 2961.500 | 2495.702 | 0.742 | 0.008 | 1.069 | 0.060 | 30.910 | 0.395 | OPC | LinB32 |
| *p1* | 17342.000 | 2906.874 | 1.240 | 0.324 | 2.294 | 0.173 | 12.908 | 0.568 | TIP3P | LinB32 |
| *p2* | 10894.500 | 5286.020 | 0.947 | 0.161 | 1.905 | 0.087 | 20.053 | 1.271 | TIP3P | LinB32 |
| *p3* | 5389.500 | 1139.447 | 0.776 | 0.021 | 1.369 | 0.115 | 19.528 | 1.153 | TIP3P | LinB32 |
| *ST* | 3066.750 | 1298.469 | 0.734 | 0.008 | 0.999 | 0.061 | 31.731 | 2.185 | TIP3P | LinB32 |
| *p1* | 14784.000 | 3342.503 | 1.018 | 0.265 | 1.937 | 0.470 | 13.206 | 1.178 | OPC | LinB86 |
| *p2* | 12679.750 | 1845.948 | 1.055 | 0.130 | 1.984 | 0.083 | 19.411 | 0.588 | OPC | LinB86 |
| *p3* | 11834.500 | 3261.131 | 0.867 | 0.060 | 1.690 | 0.277 | 16.158 | 0.286 | OPC | LinB86 |
| *ST* | 1435.000 | 1033.531 | 0.728 | 0.004 | 1.020 | 0.160 | 34.093 | 1.155 | OPC | LinB86 |
| *p1* | 15242.500 | 2775.916 | 1.132 | 0.317 | 2.281 | 0.275 | 13.086 | 1.208 | TIP3P | LinB86 |
| *p2* | 8716.750 | 2963.842 | 1.001 | 0.065 | 1.978 | 0.142 | 19.911 | 1.043 | TIP3P | LinB86 |
| *p3* | 13637.750 | 951.736 | 0.871 | 0.040 | 1.890 | 0.157 | 16.178 | 0.109 | TIP3P | LinB86 |
| *ST* | 2972.000 | 525.176 | 0.736 | 0.006 | 1.085 | 0.119 | 32.670 | 0.988 | TIP3P | LinB86 |

**Table S3B. Results of two sample t-test performed for data represented in Figure 2 for LinBWT.**

| ***Tunnel*** | ***P1*** | | ***P2*** | | ***P3*** | | ***ST*** | |
|---|---|---|---|---|---|---|---|---|
| ***Metric*** | **t-statistic** | **p-value** | **t-statistic** | **p-value** | **t-statistic** | **p-value** | **t-statistic** | **p-value** |
| *Avg_Frames* | 2.3122 | 0.0818 | -0.6999 | 0.5226 | -0.8552 | 0.4406 | -1.6735 | 0.1695 |
| *Avg_BR* | 0.0694 | 0.9480 | NA | NA | 1.6309 | 0.1782 | 0.9286 | 0.4216 |
| *Max_BR* | 1.8512 | 0.1378 | NA | NA | -0.1395 | 0.8958 | 1.2173 | 0.3105 |
| *Avg_Length* | -2.6600 | 0.0564 | NA | NA | -0.4046 | 0.7065 | 0.0668 | 0.9509 |

**Table S3C. Results of two sample t-test performed for data represented in Figure 2 for LinB32.**

| ***Tunnel*** | ***P1*** | | ***P2*** | | ***P3*** | | ***ST*** | |
|---|---|---|---|---|---|---|---|---|
| ***Metric*** | **t-statistic** | **p-value** | **t-statistic** | **p-value** | **t-statistic** | **p-value** | **t-statistic** | **p-value** |
| *Avg_Frames* | -0.5261 | 0.6177 | -0.0268 | 0.9795 | -1.5185 | 0.1797 | -0.0748 | 0.9428 |
| *Avg_BR* | -0.1513 | 0.8847 | 0.9023 | 0.4017 | -0.4351 | 0.6816 | -0.3948 | 0.7131 |
| *Max_BR* | -1.0163 | 0.3487 | 0.9889 | 0.3609 | -1.0751 | 0.3314 | -0.2517 | 0.8137 |
| *Avg_Length* | -0.2663 | 0.7989 | -0.1028 | 0.9214 | -0.2246 | 0.8312 | -0.2639 | 0.8049 |

**Table S3D. Results of two sample t-test performed for data represented in Figure 2 for LinB86.**

| ***Tunnel*** | ***P1*** | | ***P2*** | | ***P3*** | | ***ST*** | |
|---|---|---|---|---|---|---|---|---|
| ***Metric*** | **t-statistic** | **p-value** | **t-statistic** | **p-value** | **t-statistic** | **p-value** | **t-statistic** | **p-value** |
| *Avg_Frames* | -0.2111 | 0.8398 | 2.2700 | 0.0637 | -1.0616 | 0.3293 | -2.6516 | 0.0379 |
| *Avg_BR* | -0.5558 | 0.5985 | 0.7585 | 0.4769 | -0.0907 | 0.9307 | 0.3021 | 0.7747 |
| *Max_BR* | -1.2661 | 0.2524 | 0.0757 | 0.9421 | -1.2546 | 0.2563 | -0.4591 | 0.6654 |
| *Avg_Length* | 0.1422 | 0.8916 | -0.8365 | 0.4349 | -0.1291 | 0.9015 | -1.7157 | 0.1469 |

**Welch t-test performed revealed statistically significantt differences for the Avg_BR of LinBWT and LinB32 in the TIP3P model (p = 0.0428), while the OPC comparison was not significant (p = 0.0762). For LinBWT and LinB86, both OPC (p = 0.0104) and TIP3P (p = 0.0256) showed statistically significant differences for Avg_BR.*

**Table S4. Results of two sample t-test performed for data represented in Figure 3.**

| **Enzyme** | ***LinBWT*** | | **LinB32** | | **LinB86** | |
|---|---|---|---|---|---|---|
| ***Bin Center [Å]*** | **t-statistic** | **p-value** | **t-statistic** | **p-value** | **t-statistic** | **p-value** |
| *0.70* | -1.3868 | 0.2378 | -0.3743 | 0.7211 | -0.3498 | 0.7384 |
| *0.80* | -1.4980 | 0.2085 | -0.3280 | 0.7540 | -0.4019 | 0.7017 |
| *0.90* | -1.4130 | 0.2305 | -0.1110 | 0.9152 | -0.6608 | 0.5333 |
| *1.00* | -1.4104 | 0.2312 | 0.3089 | 0.7679 | -0.4336 | 0.6798 |
| *1.10* | -1.5152 | 0.2043 | 1.5031 | 0.1835 | -0.1671 | 0.8728 |
| *1.20* | -0.8944 | 0.4217 | 2.4419 | 0.0503 | 0.0280 | 0.9786 |
| *1.30* | -0.9993 | 0.3742 | 1.6144 | 0.1576 | 0.0894 | 0.9316 |
| *1.40* | -1.1590 | 0.3109 | 1.1939 | 0.2776 | 0.1518 | 0.8844 |
| *1.50* | -0.9991 | 0.3743 | 0.7589 | 0.4767 | 0.1980 | 0.8496 |
| *1.60* | -1.1575 | 0.3115 | 0.3962 | 0.7057 | 0.3898 | 0.7102 |
| *1.70* | -1.3545 | 0.2470 | 0.1118 | 0.9146 | 0.5248 | 0.6185 |
| *1.80* | -1.4336 | 0.2250 | -0.1609 | 0.8774 | 0.7840 | 0.4629 |
| *1.90* | -1.3633 | 0.2445 | -0.4236 | 0.6866 | 1.0395 | 0.3386 |
| *2.00* | -1.3166 | 0.2583 | -0.5900 | 0.5767 | 1.1446 | 0.2960 |
| *2.10* | -1.8801 | 0.1333 | -0.7543 | 0.4792 | 1.2267 | 0.2659 |
| *2.20* | -1.8924 | 0.1314 | -0.9520 | 0.3779 | 1.3851 | 0.2153 |
| *2.30* | -1.7320 | 0.1583 | -1.0410 | 0.3380 | 1.5987 | 0.1610 |
| *2.40* | -1.6594 | 0.1724 | -1.0912 | 0.3170 | 1.4280 | 0.2032 |
| *2.50* | -2.0642 | 0.1079 | -1.0000 | 0.3559 | 1.0000 | 0.3559 |

**Table S5. Results of two sample t-test performed for data represented in Figure 4**

| Enzyme | LinBWT | | LinB32 | | LinB86 | |
|---|---|---|---|---|---|---|
| ***Bin Center [Å]*** | *t-statistic* | *p-value* | *t-statistic* | *p-value* | *t-statistic* | *p-value* |
| *0.02* | -3.5402 | 0.0240 | -1.0397 | 0.3386 | -1.3760 | 0.2180 |
| *0.05* | -2.4417 | 0.0711 | 2.3849 | 0.0544 | 1.6077 | 0.1590 |
| *0.10* | -0.3786 | 0.7242 | 1.1348 | 0.2998 | 1.0981 | 0.3142 |
| *0.15* | 1.9792 | 0.1189 | 0.9270 | 0.3897 | 1.9546 | 0.0984 |
| *0.20* | 0.1347 | 0.8994 | 0.1938 | 0.8527 | 1.0028 | 0.3547 |
| *0.25* | 1.1355 | 0.3196 | 0.3500 | 0.7383 | 0.4669 | 0.6570 |
| *0.30* | 0.9456 | 0.3979 | -0.1196 | 0.9087 | 0.9146 | 0.3957 |
| *0.35* | 3.2082 | 0.0326 | -0.5130 | 0.6263 | 0.2465 | 0.8135 |
| *0.40* | 1.5285 | 0.2011 | -0.4606 | 0.6613 | 0.6680 | 0.5290 |
| *0.45* | 2.0862 | 0.1053 | -0.2981 | 0.7757 | 0.2736 | 0.7936 |
| *0.50* | 0.6917 | 0.5272 | -0.6641 | 0.5313 | 0.3453 | 0.7416 |
| *0.55* | 2.0461 | 0.1102 | -0.6795 | 0.5221 | 0.2614 | 0.8025 |
| *0.60* | 1.4816 | 0.2126 | -0.4891 | 0.6422 | 0.4847 | 0.6451 |
| *0.65* | 0.9909 | 0.3778 | -1.1438 | 0.2963 | -0.5522 | 0.6008 |
| *0.70* | 2.8016 | 0.0487 | -0.5008 | 0.6343 | -0.4713 | 0.6541 |
| *0.75* | 1.3753 | 0.2410 | -1.2426 | 0.2604 | 0.3920 | 0.7086 |
| *0.80* | 1.1548 | 0.3125 | -1.3377 | 0.2295 | 1.2125 | 0.2709 |
| *0.85* | 2.6415 | 0.0575 | -0.4570 | 0.6637 | 0.3813 | 0.7161 |
| *0.90* | 2.0207 | 0.1134 | -1.0705 | 0.3256 | -0.1675 | 0.8725 |
| *0.95* | 1.1436 | 0.3166 | -1.1600 | 0.2901 | -1.2732 | 0.2501 |
| *1.00* | 1.9181 | 0.1276 | -1.2436 | 0.2600 | -0.0539 | 0.9587 |
| *2.00* | -0.1836 | 0.8632 | -1.1945 | 0.2774 | 0.6951 | 0.5130 |
| *4.00* | -0.8623 | 0.4372 | -1.0945 | 0.3157 | 1.5445 | 0.1734 |
| *6.00* | -1.5519 | 0.1956 | -1.0000 | 0.3559 | NA | NA |
| *8.00* | 0.0534 | 0.9600 | -1.0000 | 0.3559 | NA | NA |
